# pUL36 de-ubiquitinase activity augments both the initiation and progression of lytic virus infection in IFN–primed cells

**DOI:** 10.1101/2021.01.29.428770

**Authors:** Jonas Mohnke, Irmgard Stark, Mara Fischer, Arnhild Grothey, Peter O’Hare, Beate Sodeik, Florian Erhard, Lars Dölken, Thomas Hennig

**Author notes:** **Please send correspondence to:** Dr. Thomas Hennig. contributed equally to the work (Jonas Mohnke, Irmgard Stark).

## Abstract

The conserved, structural HSV-1 tegument protein pUL36 is essential for both virus entry and assembly. While its N-terminal de-ubiquitinase (DUB) activity is dispensable for infection in cell culture, it is required for efficient virus spread *in vivo* by acting as a potent viral immune evasin. Here, we show that the pUL36 DUB activity was required to overcome interferon-(IFN)-mediated suppression of both plaque initiation and progression to productive infection. Immediately upon virus entry, incoming tegument-derived pUL36-DUB activity helped the virus to escape intrinsic antiviral resistance and efficiently initiate lytic virus replication in IFN-primed cells. Subsequently, *de novo* expressed pUL36-DUB augmented the efficiency of productive infection and virus yield. Interestingly, removal of IFN shortly after inoculation only resulted in a partial rescue of plaque formation, indicating that an IFN-induced defense mechanism eliminates invading virus particles unless counteracted by pUL36-DUB activity. Taken together, we demonstrated that the pUL36 DUB disarms IFN-induced antiviral responses at two levels, namely, to protect the infectivity of invading virus as well as to augment productive virus replication in IFN-primed cells.

**Author Summary:** HSV-1 is an ubiquitous human pathogen that is responsible for common cold sores but may also cause life-threatening disease. pUL36 is an essential and conserved protein of infectious herpesvirus virions with a unique de-ubiquitinating (DUB) activity. The pUL36 DUB is dispensable for efficient virus infection in cell culture but represents an important viral immune evasin *in vivo*. Here, we showed that tegument-derived DUB activity delivered by the invading virus particles is required to overcome IFN-induced host resistance and to initiate efficient lytic infection. *De novo* expressed pUL36 DUB subsequently augments productive infection and virus yield. These data indicate that the pUL36 DUB antagonizes the activity of yet unidentified IFN-inducible E3 ligases to facilitate productive infection at multiple levels. Our findings underscore the therapeutic potential of targeting conserved herpesvirus DUBs to prevent or treat herpesvirus disease.

## Introduction

To limit and suppress viral infection, cells have evolved a rich arsenal of restriction factors, which collectively form the innate antiviral immune system. Inherent mechanisms that operate in cells without stimulation, referred to as intrinsic immunity, include factors that prevent (e.g. TRIM-5α), sense (e.g. RIG-I, TLR3), or suppress the progression of infection (e.g. PML or PKR). The expression levels of intrinsic effectors substantially vary between different cell types and tissues but can be stimulated by the activity of interferons (IFNs) (1). Such genes are referred to as IFN-stimulated genes (ISGs). When cells sense infection via recognition of viral pathogen-associated molecular patterns (PAMPs) through pattern recognition receptors (PRRs), they activate signaling cascades culminating in the IRF3- and NF-κB-mediated induction and secretion of type I IFNs (1–3). Through autocrine and paracrine activation of the type I IFN receptor (IFNAR), the expression of several hundred ISGs is stimulated (4–6). The production of ISGs in turn installs an antiviral state in both infected as well as neighboring, uninfected cells, which subsequently hampers both susceptibility as well as permissiveness to productive infection.

Herpes simplex virus type 1 (HSV-1) is a double-stranded DNA virus that productively infects a large variety of cell types. IFN-induced effector mechanisms inhibit HSV-1 at various stages of its life cycle (7, 8) including cell entry (9), initiation of IE gene expression (10–18) and efficient viral protein production (19). To counteract these antiviral effectors, HSV-1 has evolved a plethora of viral immune evasins that efficiently interfere with the innate immune system during lytic infection (reviewed in (20)). Many of the host’s innate defense mechanisms depend on cycles of ubiquitination/de-ubiquitination of target proteins to modulate their function (reviewed in (21, 22)). In turn, HSV-1 usurps the ubiquitin system to counter these defenses through the functions of the viral proteins ICP0 (reviewed in (23)) and pUL36 (see below).

Herein, we characterized the function of the essential large tegument protein pUL36 (also named VP1-2) in the context of an established innate immune response. pUL36 is well-conserved among the different herpesviruses and contributes multiple indispensable functions both early (tegument-associated) and late (free and tegument-associated) in the herpesvirus life cycle (24–29). Upon entry of a virus particle into a cell, pUL36 is retained on the incoming capsid to facilitate microtubule-mediated capsid transport towards the nucleus and docking at the nuclear pores (24, 26, 29–36). Late in infection, the pUL36 protein is essential for secondary envelopment and tegument assembly (24, 27, 29, 34). Besides its essential, structural functions, pUL36 and its homologs also comprise an N-terminal de-ubiquitinating enzymatic activity (DUB) (37, 38) which, despite its conservation, is not essential for HSV-1 replication *in vitro* (39). Interestingly, an N-terminal pUL36 fragment, which comprises the catalytically-active DUB domain, can be detected in infected cells at late times of infection (38). On the molecular level, the DUB is a cysteine protease that can cleave both K48- and K63-linked poly-ubiquitin chains (38, 39). As such DUB activity maintains pUL36 stability by de-ubiquitinating itself (39). In other herpesviruses, the DUB homologs fulfil a variety of different functions ranging from antagonizing the innate immune and DNA damage responses, facilitating appropriate intracellular capsid motility and tegument assembly, and augmenting infection both *in vitro* and *in vivo* (40–51).

Since ubiquitination/de-ubiquitination cycles are integral to innate immunity signaling, the HSV-1 pUL36 DUB might also interfere with innate immunity at multiple levels: (i) IFN induction, (ii) IFN signaling (after IFNAR activation) and (iii) ISG functions. Indeed, the pUL36 DUB actively contributes to reducing the induction of type I IFNs by reversing the ubiquitination of TRAF3 (49) and STING (50), and pUL36 inhibits type I IFN signaling by binding to the IFNAR independently of its DUB activity (52).

However, its functions during an established IFN response have not been characterized so far. Herein, we show that the HSV-1 pUL36 DUB is required to disarm IFN-induced antiviral effector mechanisms that interfere with the induction of lytic infection at single cell level. These effects were mediated by tegument-associated DUB activity. Moreover, we demonstrate that the DUB was also required later in infection for efficient production of progeny virus in IFN-treated cells. Collectively, our data highlight an important role of the pUL36 DUB in antagonizing the IFN-induced and ubiquitination-mediated cellular defense against HSV-1 both very early, in its tegument-associated form, and late in infection.

## Results

### Comparison of replication characteristics of parental and pUL36.C65A HSV-1 strains

Despite its importance for HSV-1 spread in animal models, the DUB of pUL36 contributes little to productive infection in cell culture (39, 50). We thus hypothesized that it might augment/maintain viral replication in the presence of pre-established or ongoing innate immune responses as this induces the expression of many host proteins involved in ubiquitin-mediated pathways (Interferome database v2.0 (53)). We thus tested whether pre-treatment with type I or type I combined with type II interferons affected multi-cycle growth kinetics of the catalytically inactive DUB mutant HSV1-(17^+^)Lox-CheVP26-UL36.C65A (mCh-VP26 DUB mutant) in comparison to its parental strain. As previously reported (39, 50), there was little difference in viral yield from different cell lines among both viruses in the absence of IFN (Fig. 1A and S1). Consistent with previous reports (8, 54), there was a mild reduction in virus yield of parental virus in IFN-α treated Vero cells (Fig. 1A). A stronger inhibition was attained by combining type I and II IFN (Fig. 1A) as reported earlier (7). Of note, upon combined IFN-α and-γ pre-treatment, we had to use a 100-fold higher dose of virus to observe any CPE by 5 dpi with the parental HSV-1 strain. An enhanced inhibitory effect of the IFN pre-treatment regimens on the DUB mutant was already apparent prior to harvesting since a lesser CPE was observed compared to the parental virus. This translated into a more pronounced reduction of virus yield of the DUB mutant compared to the parental strain as early as 24 hpi. At later time points, IFN-α pre-treatment alone reduced virus yield of the DUB mutant by 10-fold, which increased to over 100-fold upon combined IFN treatment (Fig. 1A). Accordingly, we observed with IFN-α alone a 30%, and with IFN-α in combination with IFN-γ a 70% reduction in plaque size for the DUB mutant in comparison to its parental HSV-1 KOS strain (Fig. 1B). Of note, we also noted a modest, but nevertheless reproducible reduction in plaque numbers for the DUB mutants compared to their respective parental strains (c.f. below).

**Figure 1:**
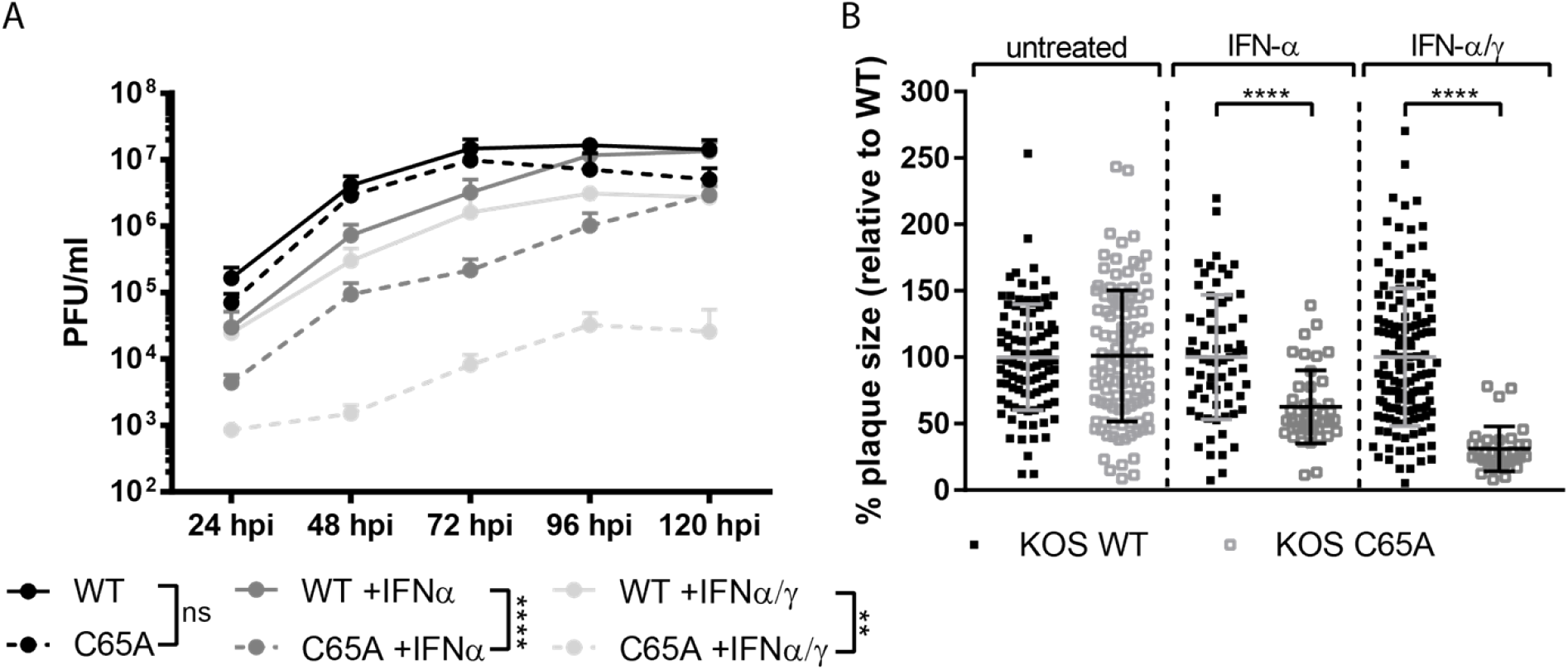
Growth kinetics of parental HSV-1 and its pUL36 DUB mutants. (A) Multi-step growth curves from untreated (black line), 500 IU/ml IFN-α-treated (dark grey) or 500 IU/ml IFN-α- and 100 IU/ml IFN-γ-treated Vero cells (light grey) (n=3). Cells were infected with an MOI of 0.001 (untreated and IFN-α) or 0.1 (IFN-α/γ) of either parental HSV-1 expressing VP26-mCherry or isogenic DUB mutant (C65A) viruses. IFN was added 20 h prior to inoculation and replenished after inoculation. Total virus yield was determined in 24 h intervals up to 120 hpi by plaque assay on IFN-naive Vero cells. (B) Plaque size from KOS parental virus or DUB mutant infected Vero cells: results from a representative experiments are shown. Untreated or IFN-treated (as in A) Vero cells were infected with ~100 PFU/ml (100-fold more for IFN-α/γ) and fixed at 2 dpi for untreated cells and at 4 dpi for IFN-treated cells. Plaque size was measured using Fiji software. (A, B) Statistical analysis was performed using the Two-Way ANOVA analysis plug-in of Graphpad Prism. ns – not significant, ** p < 0.01, **** p < 0.0001

We then asked whether the drop in yield after multi-cycle replication and the associated reduction in plaque size of the DUB mutant were due to a reduced production of infectious virions per cell, or due to a delay in productive infection. We thus performed high MOI infection, and analyzed the yield at 24 hpi from Vero cells using an MOI of 10 for untreated cells, an MOI of 20 for IFN-α, and an MOI of 100 for cells pre-treated with both IFN-α and IFN-γ (Fig S2). Microscopic pre-analyses had indicated that these virus doses ensure that all cells were synchronously infected. Even in the absence of IFN treatment, there was already a modest reduction in yield by 2.8-fold for the DUB mutant compared to parental virus. While IFN-α had little effect on the parental virus, it lead to a further 1.4-fold greater drop in yield for the DUB mutant. Furthermore, the combined pre-treatment with IFN-α and IFN-γ reduced the titer of the parental virus by 3.2-fold, but of the DUB mutant by 9.4-fold. Collectively, these data indicate that the pUL36 DUB activity is required for efficient progeny production in IFN-pre-treated Vero cells. This explains the strong replication defect of HSV-1 mutants lacking the DUB activity in multi-step growth curves.

### DUB activity is required for plaque initiation in IFN-pre-treated cells

We next asked whether the tegument-associated pUL36 DUB activity plays any role in disarming IFN-induced host defense mechanisms that act immediately upon cell entry, and thereby affect specific infectivity or host susceptibility. To test this, we analyzed the efficiency of plaque initiation, i.e. the number of induced virus plaques, for the parental and DUB mutant viruses in IFN-α-treated and untreated cells. We divided the number of plaques in IFN induced cultures by that from untreated cells to express the effect of IFN treatment in a relative plaque ratio. In these experiments, we compared the pUL36 DUB mutants of HSV1(17^+^)Lox-CheVP26 (generated here and used in (50)) and HSV1 (KOS) (described earlier (39)). For both HSV-1 parental strains, the IFN-α induction with 500 IU/ml decreased the number of plaques by about 1.5-to 2-fold (Fig. 2A), or in other words, nearly doubled the particle-to-pfu ratio, as both the mock and the IFN-treated cells were inoculated with the same virus dose. Moreover, more extensive IFN treatment (500 IU/ml IFN-α + 100 IU/ml IFN-γ vs. 500 IU/ml IFN-α vs. 100 IU/ml IFN-α) lead to a stronger repression of plaque formation (Fig. S3). Importantly, IFN-induction reduced plaque formation of both HSV-1 strain (17^+^ and KOS) DUB mutants significantly stronger than that of their respective parental strains (p < 0.01 for both DUB mutants), consistent with a further increase in the particle-to-pfu ratio by about two-fold (Fig. 2A and S3A, C). This effect was evident both at low MOI (i.e. 100-200 PFU per 6-well) and at high MOI using serially diluted inocula (Fig. S4). Immunolabeling for the immediate early HSV-1 protein ICP4 at 24 hpi ensured that even very small foci and plaques consisting of few infected cells were not missed.

**Figure 2:**
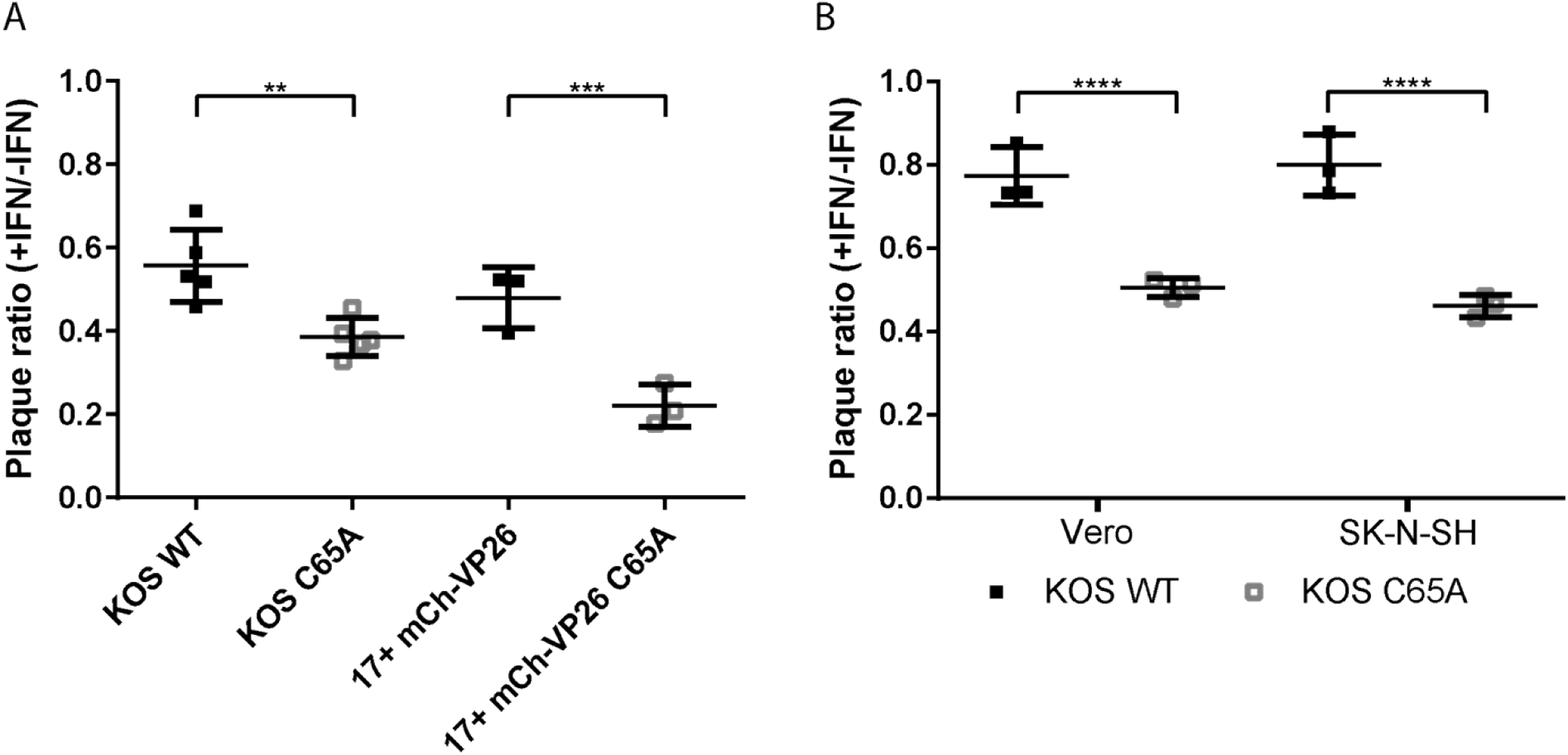
Effect of the pUL36 DUB on plaque initiation. Vero cells were treated with 500 IU/ml IFN-α or left untreated. Plaque assays were performed 20 h after treatment of cells with IFN using ~100 PFU/well of either parental HSV-1 (strain KOS or the mCherry-VP26 derivative of stain 17) or their isogenic DUB mutants. The ratio of plaques that formed in presence and absence of IFN is shown. Using the same experimental setup as in (A), comparable results were observed for KOS parental virus and the isogenic DUB mutant in both Vero and human SK-N-SH cells. (A, B) Statistical analysis was performed using the One-Way ANOVA analysis plug-in of Graphpad Prism.

While we had initially analyzed the pUL36-DUB mutants in Vero cells, we next asked whether they were also impaired in the initiation of plaque formation in human SK-N-SH, RPE-1, and HaCaT cells. Consistent with our previous results, the DUB mutants generated significantly fewer plaques in human cells after pre-treatment with IFN-α (Fig. 2B, S3B). These data highlight the role of the pUL36 DUB activity in augmenting infectivity and suggest a defect very early in infection, indicative of a role of tegument-associated pUL36 DUB in this phenotype.

### The pUL36 DUB facilitates efficient initiation of infection independently of its effect on IFN signaling

During the course of this study, it was reported that the HSV-1 pUL36 DUB inhibits IFN signaling by binding directly to the IFN-α receptor, thereby interfering with the induction of ISG expression (*34*). As we had pre-treated our cells with IFN, it seemed unlikely that the observed reduction in plaque formation was due to an inhibition of IFN signaling by the pUL36 DUB. Nevertheless, we tested whether the DUB mutant phenotype also occurred if cells were only treated with IFN-α after inoculation with the virus. For this, Vero cells, that had been (*i*) pre-treated with 100 or 500 IU/ml IFN-α prior to inoculation only (‘pre’), (*ii*) treated throughout the whole experiment (‘pre + post’), or (*iii*) only treated with IFN-α after removal of the inoculum (‘post’), were infected with the parental KOS strain or with the DUB mutant (Fig 3A, B). Of note, the ability of the DUB mutant virus to initiate plaques was reduced to a similar extent in cells that had been treated with IFN only prior to infection compared to cells that had been treated throughout the experiment (Fig. 3A). Conversely, when IFN-α was added to the culture only after removal of the inoculum (**Error! Reference source not found.**. 3B, ‘post’), there was still a modest reduction in plaque numbers by 25% not only for the DUB mutant but also for the parental KOS virus.

**Figure 3:**
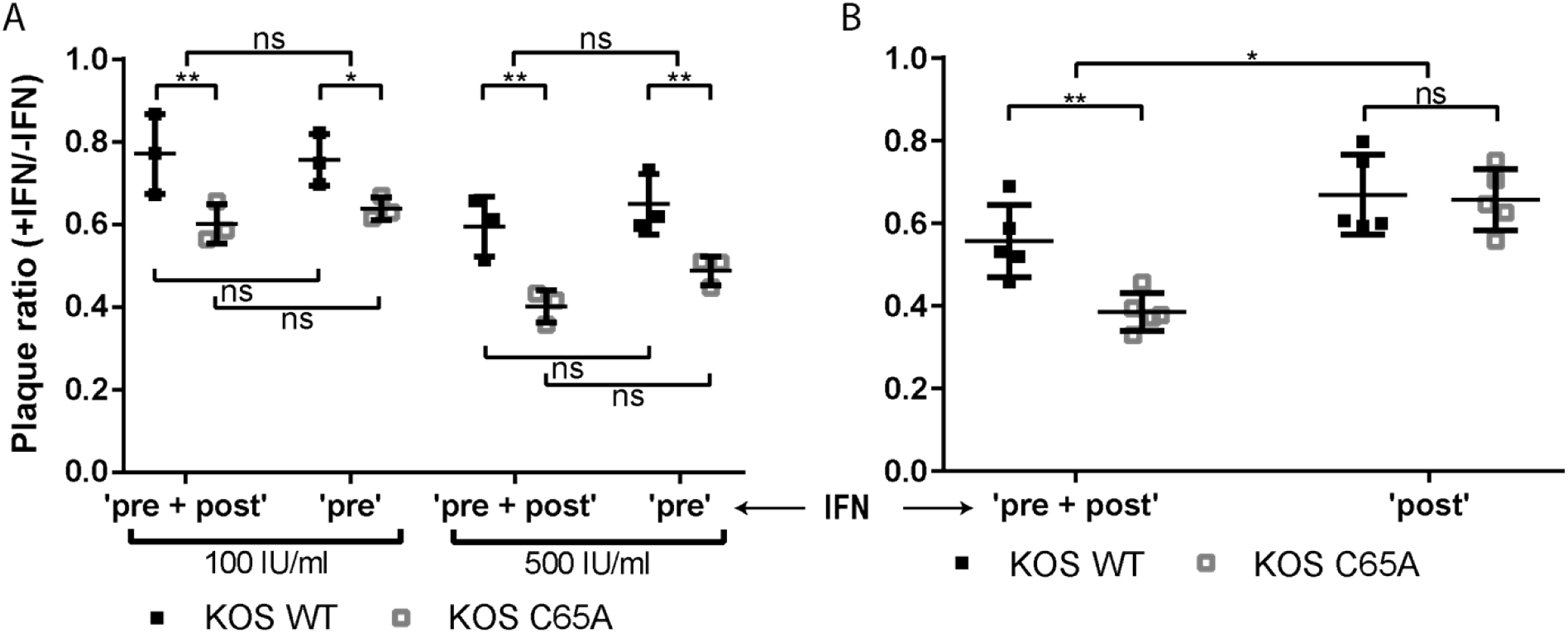
The UL36 DUB augments plaque formation in IFN pre-treated cells. (A) Vero cells, either untreated or treated with 100 or 500 IU/ml IFN-α prior to inoculation only (‘pre’) or treated with 100 or 500 IU/ml IFN-α prior to and after inoculation (‘pre + post’), were infected with ~100 PFU/well of KOS bacmid-derived parental or DUB mutant viruses. (B) As in A, but here 500 IU/ml IFN-α treatment applied throughout the whole assay (‘pre + post’) was compared to treatment with 500 IU/ml IFN-α applied only after inoculation (‘post’). (A, B) Plaque ratios from IFN-treated over untreated Vero cells were calculated and plotted. Statistical analysis was performed using the Two-Way ANOVA analysis plug-in of Graphpad Prism.

Taken together, these data indicate that HSV-1 requires the pUL36-DUB activity to efficiently initiate plaque formation in IFN pre-treated cells. In other words, the DUB disarms the anti-viral state promoted in a cell by prior IFN induction. We thus hypothesized that tegument-associated pUL36 DUB activity is required for efficient establishment of lytic infection in the very first IFN-stimulated cell of a plaque. However, the dependence on DUB activity is subsequently overrun during the progression of plaques since the cells around the initially infected cells are invaded by a much larger number of virions.

### Tegument-derived pUL36 DUB activity is required to overcome IFN-induced antiviral restriction

To test this hypothesis, we generated trans-complemented HSV-1 virions for both the parental KOS strain and its isogenic DUB mutant. The two viruses were propagated on the pUL36-expressing complementing cell line RSC-HAUL36, which efficiently rescues infectivity of pUL36 deletion mutants (27, 29, 30, 34, 55). The resulting virions (Fig. 4A, right) contain both WT pUL36 (blue), expressed from the host cell genome, and pUL36.C65A protein (red), expressed from the viral genomes. When non-complementing cells like Vero or SK-N-SH are inoculated with such virions, only virally-encoded pUL36.C65A is expressed *de novo*, thereby enabling us to assess the role of the incoming tegument-associated DUB in augmenting virion infectivity (Fig. 4A, left).

**Figure 4:**
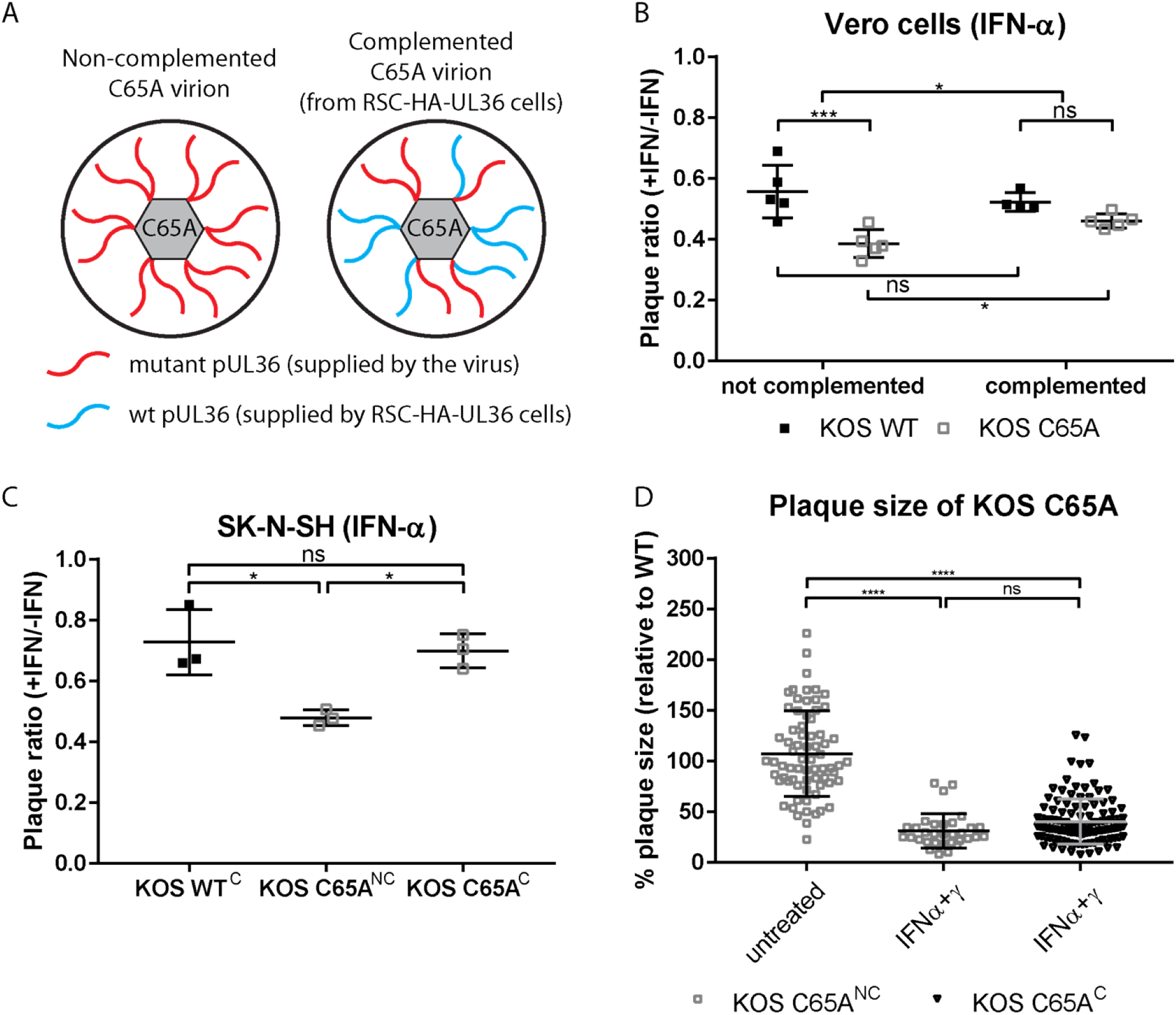
Tegument UL36 DUB activity augments plaque initiation in IFN pre-treated cells. (A) Schematic representation of non-complemented virions (left) and complemented (right) DUB mutant virions. When the DUB mutant virus (C65A) is grown on normal producer cells (not expressing UL36, e.g. RS cells) the virions, which are produced, only contain the mutant, virus-derived pUL36 DUB protein (red). When C65A virus is grown on UL36 complementing cells (e.g. RSC-HA-UL36) the resulting virions contain a mixture of mutant, virus-derived pUL36 and wild-type, cell-derived pUL36 (blue, right). Parental virus grown on the respective cells served as controls. After initial infection of non-complementing cells (e.g. Vero or SK-N-SH cells) with complemented DUB mutant, plaques progress as ‘mutant’ since virions produced in these cells will only contain the DUB mutant pUL36. (B, C) Vero or SK-N-SH cells were treated prior to and after inoculation with (B) 500 IU/ml IFN-α or (C) 25 IU/ml IFN-α and infected at ~100 PFU/well of the indicated viruses. Complementation status is indicated as “C” (complemented) or “NC” (non-complemented). Plaque ratios were calculated and plotted as before. (D) Plaque size on untreated and IFN-α/-γ pre-treated Vero cells was analyzed for complemented DUB mutant virus and compared to that of the non-complemented virus. Each data point represents one plaque which was normalized against the mean plaque size of the parental virus assayed simultaneously. (B-D) Statistical analysis was performed using the (C, D) One-way ANOVA or (B) Two-Way ANOVA analysis plug-in of Graphpad Prism.

IFN-α-or IFN-α/γ-treated Vero cells were inoculated with 100 PFU/well (100-fold more for IFN-α/γ treatment) of virions harboring only the DUB mutant pUL36 (‘not complemented’, red in Fig. 4A) or a mix of DUB mutant and wild-type pUL36 proteins (‘complemented’, red and blue in Fig. 4A) in their tegument. Interestingly, virions containing only pUL36 DUB mutant proteins generated lower plaque ratios (>50%) than those complemented with wild-type pUL36 proteins in both, IFN-α-(Fig. 4B) and IFN-α/γ-treated Vero cells (Fig. S3C). More strikingly, on SK-N-SH cells treated with 25 IU/ml IFN-α, the complemented KOS DUB mutant gave rise to plaque numbers that were indistinguishable from complemented parental virus (Fig. 4C). Notably, while the plaque ratio on Vero cells of the trans-complemented DUB mutant could be at least partially rescued, this did not increase plaque size when compared to non-complemented DUB mutant virions (Fig. 4D). We conclude that the tegument-associated pUL36 DUB enables HSV-1 to overcome IFN-induced, antiviral effector mechanisms.

### Infectivity of incoming virus is not fully rescued upon removal of IFN

We previously reported that IFN pre-treatment efficiently suppressed murine cytomegalovirus (MCMV) infection by preventing the expression of viral immediate early genes in a PML body-dependent manner (56). Upon IFN withdrawal, viral gene expression resumed and efficient lytic infection re-initiated. Since we found that IFN pre-treatment selectively reduced plaque initiation of the DUB mutant and since IFN protection against HSV-1 is known to wane within 24 h of treatment (57), IFN withdrawal at the time of inoculation should ultimately lead to the reactivation of reversibly suppressed viral genomes. To this end, we pre-treated Vero cells cultured in 96-well plates for 16 h with 500 IU/ml of IFN-α We then infected them either with HSV1(17^+^)-CheVP26 or its respective DUB mutant at 1.5 PFU/well for six days in presence (“pre + post”) or absence (“pre only”) of IFN-α. An infection with 1.5 PFU/well should give rise to about 77% infected wells on average. At 6 dpi, the supernatants were transferred to untreated Vero cells that were then cultured for four days to detect with high sensitivity any infectious virus that was released from the inoculated cells of the first round. Although we had co-titrated the parental and DUB mutant virus, we observed between 70 to 80 infected wells for the parental HSV-1 strain, but slightly higher numbers for the DUB mutant. This could be converted to an inoculation dose of 1.5 PFU/well for parental and 2.6 PFU/well for the DUB mutant virus (Fig. 5B). Nevertheless, and consistent with the plaque reduction assay, continuous IFN-α-treatment (“pre + post”) reduced the infectivity of the parental by only ~2-fold to 0.7 PFU/well but of the DUB mutant by 4.4 fold from 2.6 to 0.6 PFU/well. Interestingly, neither the infectivity of the parental nor of the DUB mutant fully recovered upon IFN-α withdrawal at the time of inoculation. Recovery of the DUB mutant was only marginally greater than for its parental strain despite the significantly stronger IFN-α-mediated suppression. These findings are consistent with the hypothesis that IFN induces distinct antiviral effector mechanisms that either reversibly or irreversibly suppress invading HSV-1 capsids or genomes. These are disarmed by the tegument-associated pUL36 DUB activity. The underlying molecular mechanisms remain to be elucidated.

**Figure 5:**
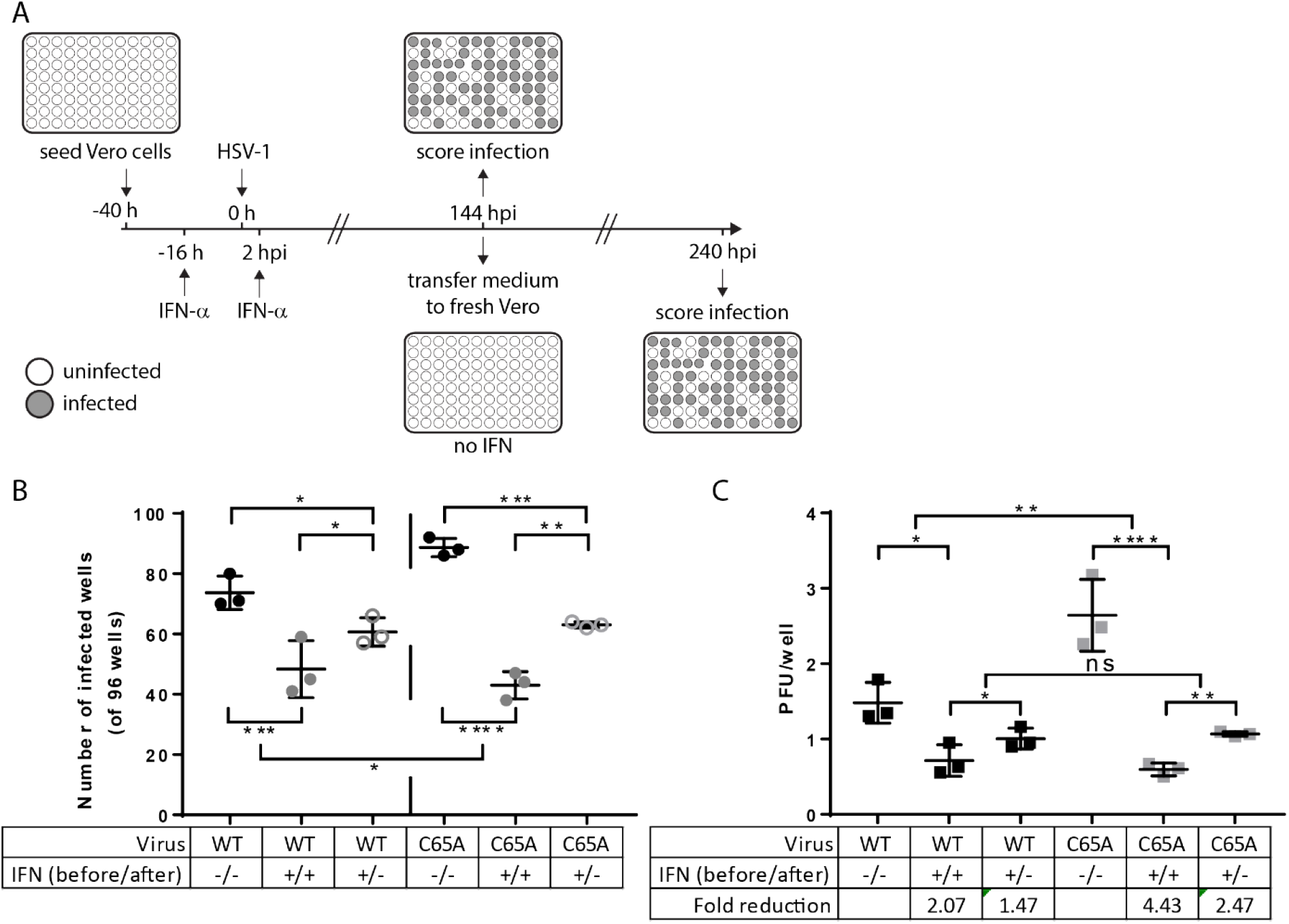
IFN-mediated inhibition of plaque initiation is partially irreversibly and greater for the C65A mutant. (A) Schematic of the experimental conditions: Vero cells were either left untreated (−/−) or were treated with 500 IU/ml IFN-α both prior to and after infection (+/+). To check for reversibility of the IFN-induced inhibition of plaque initiation, we now also included a third condition in which IFN was removed from the infected cells immediately prior to virus inoculation (+/−). Cells were infected with approximately 2 PFU/well of mCh-VP26 bacmid-derived parental or DUB mutant viruses. At six days post infection infected wells were scored by fluorescence microscopy (mCherry, indicated as grey wells). Additionally, supernatant from each well was transferred to IFN-naïve Vero cells to ensure the detection of even low levels of productive virus replication. After a further four days, infected wells (mCherry) were scored using fluorescence microscopy and finally by crystal violet staining. (B) Number of infected wells per 96-well plate from the three independent experiments are shown. Only scores obtained after the full ten days are shown. (C) Based on the percentage of infected wells, the infectivity of the virus inoculum (in PFU/well) under the respective treatment conditions was determined using a Poisson distribution and plotted for each virus and treatment condition. (B, C) Statistical analysis was performed using Two-Way ANOVA analysis plug-in of Graphpad Prism.

## Discussion

Although pUL36 performs essential functions during HSV-1 infection, the role of the conserved DUB domain remain poorly understood. Herein, we demonstrate that the DUB activity exerts at least two functions during HSV-1 infection of IFN-stimulated cells; it augments both virus infectivity likely prior to nuclear targeting of capsids and the production of infectious progeny.

The reduced infectivity of the DUB mutants could be explained by several scenarios. Capsids lacking tegument pUL36 DUB activity might be more susceptible to cytosolic ubiquitin-dependent degradation, or they might be impaired in their targeting to the nuclear pores. Alternatively, the genomes of DUB mutants might be repressed more efficiently upon entry into the nucleus.

Studies on pseudorabies virus, an alphaherpesvirus of swine, indicate that the tegument-associated DUB modulates ubiquitination of incoming capsids (on pUL36 itself) during axonal transport of incoming capsids to nuclei of neurons (51, 58). Moreover, herpesvirus capsid stability during entry appears to be a critical factor for appropriate uncoating at the nuclear pore, as incoming capsids may be targeted for proteasomal degradation in the cytoplasm (47, 59) or by the IFN-inducible GTPase MxB (9, 60). However, no evidence has been reported that the latter involves ubiquitination. Taken together with the finding that MG-132 inhibits capsid motility towards the nucleus (61, 62), this is consistent with a role of ubiquitin during virus entry (reviewed in (63)). However, there is conflicting evidence regarding the activity of MG-132 during early HSV-1 infection (64). We propose that one or more IFN-induced ubiquitin E3 ligase(s) restrict incoming capsids by mis-routing or inhibiting their microtubule transport or by virion degradation. Apparently, alphaherpesviruses learned to reverse and disarm this inhibition through the activity of their tegument-associated DUB enzyme. This is consistent with our findings that ‘pre-’ but not ‘post-‘ IFN treatment restricted plaque initiation of the DUB mutant and that this was reversed when the virions were trans-complemented with the active pUL36 DUB enzyme.

While Horan et al. show that the major capsid protein VP5 is ubiquitinated in macrophages (59), the viral protein targets of the IFN-induced ubiquitination remain to be elucidated. Notably, upon infection with a HSV-1 pUL36 DUB mutant, the viral structural proteins pUL36, pUL37, pUL25, pUL6, and VP5 show a higher level of ubiquitination in DUB mutant-infected cells compared with the corresponding parental strain (58). In this respect, pUL36 itself might be the most relevant substrate as it (*i*) contains at least four ubiquitination sites (58), (*ii*) is ubiquitinated during cell entry (58), and (*iii*) possesses auto-catalytic DUB activity (39).

Several studies suggested that HSV-1 genomes injected into the nucleus of IFN pre-treated cells cannot initiate transcription (65, 66). This restriction is exacerbated in HSV-1 mutants lacking the viral E3 RING ubiquitin ligase ICP0 (54, 67–69). We previously reported that treatment with IFN-β reversibly silenced MCMV infection in endothelial and fibroblast cells (56). Incoming viral genomes of the pUL36 DUB mutants might be silenced through the activity of nuclear host E3 ligases such as RNF8 as suggested for ICP0-null virus (70, 71). The pUL36 DUB, after being cleaved at the nuclear pores from the capsids (72), might be imported into the nucleoplasm, and remove ubiquitin from these targeted host proteins to relieve this repression. In contrast to MCMV, whose infectivity was completely rescued upon IFN withdrawal (56), IFN-induced reduction of HSV-1 infectivity was only reversed by about 50%. These data indicate that additional, non-reversible IFN-induced defense mechanisms repress HSV-1 infection. More importantly, since there was a similar recovery for the parental virus and UL36 DUB mutant, the DUB apparently does not appear to counteract this irreversible IFN-induced defense mechanism. We thus favor a model where the observed effects occur upstream of genome entry into the nucleus.

## Materials and methods

### Cells

Vero (Green Monkey kidney epithelial), RSC, RSC-HAUL36 (27), retinal pigment epithelial (RPE)-1, SK-N-SH (human glioblastoma) and HaCaT (human keratinocytes) were cultured at 37°C and 5% CO2 in a humidified atmosphere in DMEM supplemented with 10% FBS, 1x Penicillin-Streptomycin and 1x non-essential amino acids.

### Viruses

The HSV-1 KOS-37 BAC (bacterial artificial chromosome; (KOS parental) and its derivative UL36.C65A (DUB catalytic mutant) mutant viruses are described before (39, 73). The same is true for the HSV1(17^+^)Lox-CheVP26 BAC harboring the mCherry-VP26 fusion (29). The C65A equivalent mutation in UL36 was inserted by *en passant* mutagenesis (74) using the primers listed in Table 1. Briefly, a kanamycin resistance cassette flanked by two I-SceI restriction sites was lifted from the pEP-Kan-S vector (75) using primers prW534 and prW853, and prW854 and prW855 with two consecutive PCRs. 100 ng of the gel-purified PCR product from the second PCR were transformed into GS1783 cells containing the HSV1(17^+^)Lox-CheVP26 BAC. The insertion was confirmed using restriction digestion. The kanamycin cassette was removed from correct clones by induction of the restriction enzyme *I-SceI* and *red* recombinase with 1% arabinose and 15 min incubation at 42°C, respectively. Clones were confirmed by restriction digestion and sequencing of a PCR product of prW902 and prW429 validated the introduced modifications. Furthermore, the novel BAC was subjected to NGS. The recombinant HSV-1 strains HSV1(17^+^)Lox-CheVP26and HSV1(17^+^)Lox-CheVP26-UL36.C65A (also called in HSV1(17^+^)Lox-CheVP26-pUL36ΔDUB in Bodda et al. 2020) were reconstituted by transfecting the respective BACs into BHK cells.

**Table 1.**
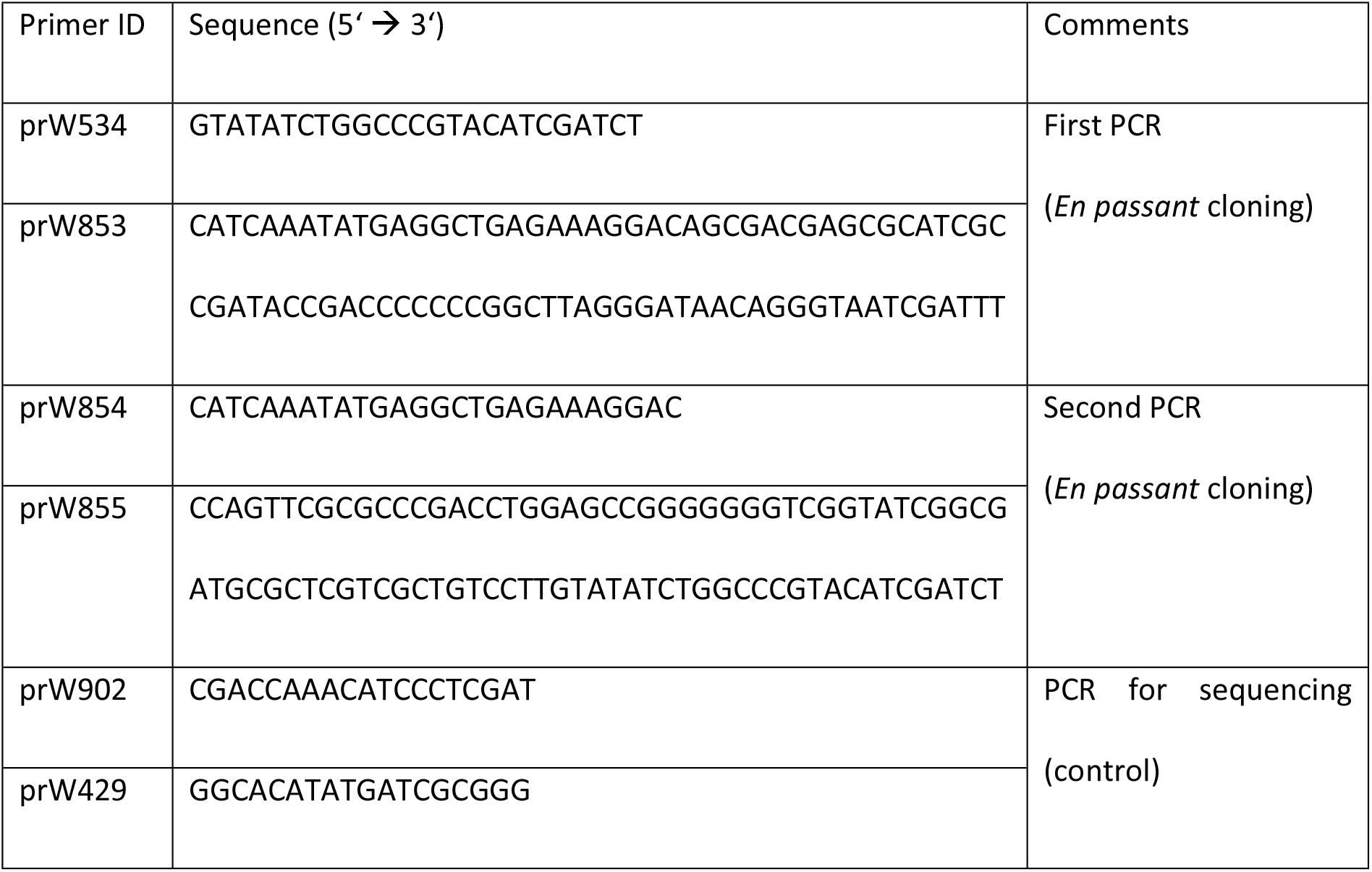
Primers used to generate HSV-1(17+)Lox-CheVP26-UL36.C65A

### Virus stock production

Viral stocks were generated in BHK or for complementation studies in RSC-HAUL36 cells. Briefly, 10 to 40 dishes (15 cm) of cells were infected at confluence using an MOI of 0.005-0.01 and maintained with 2% FBS until full cytopathic effect (CPE) was observed. Cells were harvested by scraping and pooled. Extracellular virus (ECV) and cells were separated by a 10 min centrifugation at 3000 g. The ECV was collected and cells subjected to three freeze-thaw cycles to release cell-associated virus (CAV). CAV was recovered by centrifugation (10 min, 3000g, 4°C) to remove cellular debris. Pooled ECV and CAV were concentrated at 25,000 g for 2 h and resuspended overnight in PBS on ice. The mixture was dounced 30x on ice to completely resuspend the pellet and then layered onto a continuous 5-15 % polysucrose 400 gradient. Heavy particles were separated at 25,000 g at 4°C for 2 h, collected by pipette aspiration, filled up with ice-cold PBS and concentrated at 25,000 g (4°C, 2 h), aliquoted, shock frozen in liquid nitrogen and stored at −80°C. Virus aliquots were used only once.

### Virus titration

Viruses used in this study were titrated on the cell line used for each experiment using the exact downstream assay inoculation protocol. Cells were infected at confluence and overlaid with DMEM containing 2 % FBS and 0.5 % carboxymethylcellulose. Plaques were quantified 2-4 days post-infection (dpi) after formalin fixation and crystal violet (0.1%) staining.

### Growth curves

Untreated cells were seeded 24 h prior to infection. Cells were infected at confluence for 1 h using a MOI 0.001 (multi-step) or MOI 10 (single-step yield) and maintained in DMEM containing 2 % FBS until complete CPE was observed. Total virus was harvested in 24 h intervals, and the yields were determined by plaque assays on Vero cells. Total virus was released by freeze-thawing samples three times and removing the debris by a short spin (3000 g, 10 min, 4°C).

For growth curves with IFN treatment, Vero cells (60-80 % confluent, seeded 24 h earlier), were treated with 500 IU/ml IFN-α2a (IFN-α henceforth) or with a combination of 500 IU/ml IFN-α and 100 IU/ml IFN-γ. Cells were pre-treated for 16-20 h prior to inoculation, then infected at an MOI of 0.001 (multi-step) and cultured for up to 5 dpi with samples harvested at 24 h intervals (cells scraped into medium) or infected at MOI 10 (single-step) and harvested at 24 hpi. MOIs were increased 2-fold for IFN-α and 10-fold (single step) or 100-fold (multi-step) for IFN-α/γ co-treatment. Total yields were determined using plaque assays on untreated Vero cells.

### Plaque reduction assays

Cells were seeded in 6-well plates to a density of 60-80% (24 h after seeding). Cells were treated with IFN-α (SK-N-SH 25 IU/ml, RPE-1 50 IU/ml, HaCaT 100 IU/ml, Vero 100 or 500 IU/ml) or a combination of IFN-α (500 IU/ml) and-γ (100 IU/ml) for 16-20 h prior to inoculation. A single inoculum providing 100-200 PFU to each well was prepared for all IFN-α conditions. For IFN-α/γ co-treatment inocula were serially diluted to give rise to 10^4^, 10^3^ or 10^2^ plaques. Besides these modifications, the assay functions like a standard plaque assay. Cells were inoculated for 1 h with periodic agitation. To prevent the spread of extracellular virus, cells were overlaid with 2% FBS and 0.5% carboxy-methylcellulose containing medium (+/−IFN). The cells were fixed with formalin at 2 dpi (untreated) or 4 dpi (IFN-treated), and stained with crystal violet. The plaque ratio was determined by dividing the number of plaques obtained on IFN-treated cells by the mean number of plaques obtained on the untreated cells. For the IFN-α/γ combination treatment, the opposite ratio was determined (fold inhibition of plaque formation compared to untreated).

### Determination of plaque size

Fixed and stained plaque assays of Vero cells were used to determine plaque size (from the plaque reduction assay). For each condition multiple images of multiple wells were taken at hardware pre-defined positions using the Ensight plate reader 96-well imaging plugin (n=2 for IFN-α and IFN-α/-γ). The plaque area was measured with the Fiji software package by manually outlining all visible plaques and using the measure plug-in. A minimum of 32 plaques were counted for each replicate and each plaque is represented in the corresponding graphs. All analysis was done using GraphPad Prism 7. 2-way ANOVA was used to compare the difference between parental and mutant viruses and to compare treatment pairs (interaction).

### IFN recovery assay

To confirm the results of the plaque reduction assay, we adapted the experiment to a 96-well format. A total of 6,000 Vero cells were seeded per well to obtain about 60% confluency at the start of IFN-α treatment. After 24 h, the regular growth medium was replaced by medium lacking (mock) or containing 500 IU/ml IFN-α. After 16-20 h, the media were removed, and the cells were washed with serum-free medium, and infected with 50 μl serum-free medium containing 1.5 to 3 PFU/well of HSV1(17+)Lox-CheVP26 or HSV1-CheVP26-UL36.C65A. The virus dose was calculated based on a Poisson distribution to generate a phenotypic window between the parental and DUB mutant viruses assuming a roughly two-fold drop in infectivity, which we observed in the plaque reduction assay. After 2 h, 150 μl/well of mock or IFN-α-containing media were added to yield an IFN-α concentration of 500 IU/ml. After 6 days, half of the medium was transferred to untreated Vero cells cultured in 96-well plates and incubated for another 4 days. Infected wells were scored by fluorescence microscopy (mCherry) and then after fixation and crystal violet staining. The ratio of infected wells for each condition compared to untreated was formed as before (+/−IFN-α) and plotted. Statistical testing was performed essentially as for the plaque reduction assay.

### Statistical analysis

All significance testing was done using the One- or Two-way ANOVA analysis plug-ins of GraphPad Prism 7. No corrections for multiple comparisons were conducted.

### Immunofluorescence microscopy

Vero cells were seeded in 12-well IBIDI chamber slides at 10,000 cells per well (60% confluence the next day) and treated, after 24 h, with 500 IU/ml IFN-α or a combination of 500 IU/ml IFN-α and IFN-γ for 16-20 h. Virus inocula were serially diluted ten-fold from MOI 10 to 0.01. Cells were inoculated for 1 h and then the medium was exchanged (+/−IFN). IFN-treated and untreated cells were infected with the same inoculum for each MOI. Cells were fixed at 24 hpi and stained with DAPI, anti-ICP4 (Santa Cruz) and anti-mouse Alexa Fluor 488 (Life Technologies). Infected cells were imaged using DAPI, FITC and Texas Red filter sets on a Leica DMi 8 inverted microscope.

## Acknowledgments

This work was funded by institutional funding available to TH.

## Authors’ contributions

JM, IS, MF, AG and TH performed the wet-lab work. LD and TH supervised wet-lab work. FE guided statistical analysis and calculations. LD and TH discussed and designed the experiments with the support of PO and BS. PO and BS supplied reagents to conduct the study. LD and TH analyzed the data and wrote the paper. All authors read, commented on and approved the content of the paper.

